# Scale dependency of taxonomic and functional diversity in old-growth and secondary loess steppic grasslands

**DOI:** 10.1101/2023.10.10.561714

**Authors:** Péter Török, Balázs Teleki, László Erdős, Andrea McIntosh-Buday, Eszter Ruprecht, Béla Tóthmérész

**Author notes:** The authors contributed equally to the research (Péter Török, Balázs Teleki).

## Abstract

Widespread campaigns on forest restoration and various tree planting actions lower the awareness of the importance of grasslands for carbon sequestration or biodiversity conservation. Even lower attention is given to the conservation of biodiversity and ecosystem functioning in remnants of ancient, so called old-growth grasslands. Old-growth grasslands in general harbour high biodiversity and even small patches of these can act as important refuges for many plant and animal species in urbanised or agriculture driven landscapes. Spontaneous succession of grassland is frequently viewed as a cost-effective tool for grassland restoration, but its applicability is strongly dependent on many local to landscape-scale factors and the recovery often slow. Thus, for the assessment of the effectiveness of spontaneous succession as a restoration tool, it is essential to compare the species diversity and functional attributes of old-growth and secondary grasslands. We studied the taxonomic and functional diversity in thirteen loess steppic grasslands (8 old-growth and 5 secondary) using differently sized plots ranging from 0.01 to 100m^2^. Our results indicated that there are remarkable differences in taxonomic and functional diversity between old-growth grasslands and secondary ones. We also point out that during secondary succession there is a likely functional saturation of the species assembly in the first few decades of recovery, and while patterns and structure of secondary grasslands became quite similar to those of old-growth grasslands, the species richness and diversity remains still much lower. Old-growth grasslands support considerable plant diversity, and species composition is slow to recover if destroyed for agricultural land use. This underlines the priority of protecting existing old-growth grassland remnants over restoration actions and the recovery of secondary grasslands.

## 1. Introduction

The decade between 2021-2030 is dedicated to ecosystem restoration by the United Nations, reflecting the increased global interest in ecological restoration (United Nations, 2019). Unfortunately, widespread campaigns like the Bonn Challenge or the newest EU Biodiversity strategy are highly focused on the recovery of forests by various tree planting campaigns (Tölgyesi et al., 2022). This can pose a threat to grassy habitats due to the lower awareness of the importance of grasslands for carbon sequestration or biodiversity conservation compared to forests or tree plantations, especially in arid conditions (Török et al., 2021; Buisson et al., 2022; Silveira et al., 2022). Besides grassland restoration, the conservation of biodiversity and ecosystem functioning in remnants of ancient, so called ‘old-growth’ grasslands has received even less attention.

The term ‘old-growth’ is frequently used for ‘ancient’ remnant forests of high nature value, but the application of this concept for grasslands is relatively new in the literature (Veldman et al., 2015; Nerlekar and Veldman, 2020; Buisson et al., 2022). Old-growth grasslands in general harbour high biodiversity and even small patches of these can act as important refuges for many plant and animal species in urbanised or agriculture driven landscapes (Dengler et al., 2014; Klaus, 2013; Lindborg et al., 2014). It was also stressed that the small-scale biodiversity of some of these grasslands is comparable or even might exceed the biodiversity of tropical rainforests (Wilson et al., 2012). Among temperate grasslands, natural old-growth dry grasslands are the most affected by anthropogenic activities and are highly subject to degradation (Cousins et al., 2007; Löffler et al., 2020). Old-growth dry grasslands on fertile soils, like lowland steppic grasslands are especially threatened, as their soil is the most appropriate for crop production. As a result, in most countries vast areas of steppic grasslands have been destroyed and converted to arable lands in the second half of the last century (Wesche et al., 2016; Deák et al., 2021). Steppic grasslands are valuable habitats harbouring high number of specialist species of conservation interest and several types are listed as priority habitat types in the EU Habitats Directive, thus their conservation and restoration is of high priority (e.g., 6250 - Pannonic loess steppic grasslands; European Union, 1992).

Most ecological phenomena are reported to be scale-dependent and different outcomes and patterns of their functions and biodiversity can be expected at various scales (Hewitt et al., 2017). The study of species-area relationships, starting with one of the pioneering works of McArthur-Wilson’s Island biogeography, have been in the focus of scientific interest for decades, also gaining momentum in vegetation ecology in the last decade (McArthur and Wilson, 1967; Lindgren and Cousins, 2017; Ottaviani et al., 2019). Several studies have been conducted analysing the scale-dependent patterns in vegetation composition and vegetation processes mostly focusing on relationships between biomass production, species richness, and plot size (Palmer and White, 1994; Chiarucci et al., 2006; Dembicz et al., 2020). Compared to scale-dependent patterns in species richness, the scale-dependency of some functional attributes (multi-trait functional diversity components or single-trait values) has received less attention so far (Ottaviani et al., 2019).

Spontaneous succession is frequently viewed as a cost-effective tool for grassland restoration, but its applicability is strongly dependent on the local habitat conditions and the propagule pressure from the surroundings of the restoration site, and the recovery process is strongly time-dependent, and often slow (Ruprecht, 2006; Prach et al., 2015; Török et al., 2011a). Grassland restoration is considered to be much faster compared to forest restorations because i) the development of herbaceous vegetation cover is fast and the ii) herbaceous species reach reproductive maturity earlier (Buisson et al., 2022). However, it was also indicated that the legacy effect of the subjected site, for example in form of residual soil fertility or accumulated soil seed bank of undesirable species, can also hamper the regeneration process (Török et al., 2021). It was also stressed that the community assembly of grasslands could be slow and compositional similarities between secondary grasslands and old-growth ones remain highly discrepant for several decades or even centuries (Nerlekar and Veldman, 2020; Prach et al., 2016). It was stated in former research that well-preserved old-growth grasslands are highly organised in terms of species composition and show a high degree of stability considering small-scale species fluctuations (Bartha et al., 2003; Virágh and Bartha, 2003; Teleki et al., 2020). Thus, for the assessment of the effectiveness of spontaneous succession as a restoration tool, it is essential to compare the species diversity and functional attributes of old-growth and secondary grasslands.

To analyse differences in species and functional diversity patterns and vegetation composition of secondary and old-growth grasslands, we selected altogether 13 grassland stands (8 old-growth and 5 secondary) in the South-Eastern part of Transdanubia, Hungary, Central Europe. We tested the following hypotheses: i) Species richness of secondary grasslands does not reach to levels found in old-growth grasslands // The old-growth grasslands are more diverse compared to secondary ones. In particular, we assumed that ia) species richness and Shannon diversity are lowers, ib) the dominance of the most abundant species of the plant community (Berger-Parker dominance) is higher, and ic) the functional diversity is lower in secondary grasslands compared to old-growth ones. We also hypothesised that ii) the difference in taxonomic and functional diversity between secondary and old-growth grasslands is strongly scale dependent. Finally, we analysed whether iii) there is a difference in species composition between secondary and old-growth grasslands.

## 2. Materials and methods

### 2.1. Study region

The study region is situated in the South-Eastern part of Transdanubia, Hungary, in the Tolnai-hegyhát region, near to the settlements of Szekszárd and Bonyhád (Figure 1). The climate is moderately continental with a mean annual temperature between 10-11°C and annual precipitation of 550-600 mm, characterised with a marked drought period in the mid-summer (Mezősi, 2017).

**Figure 1.**
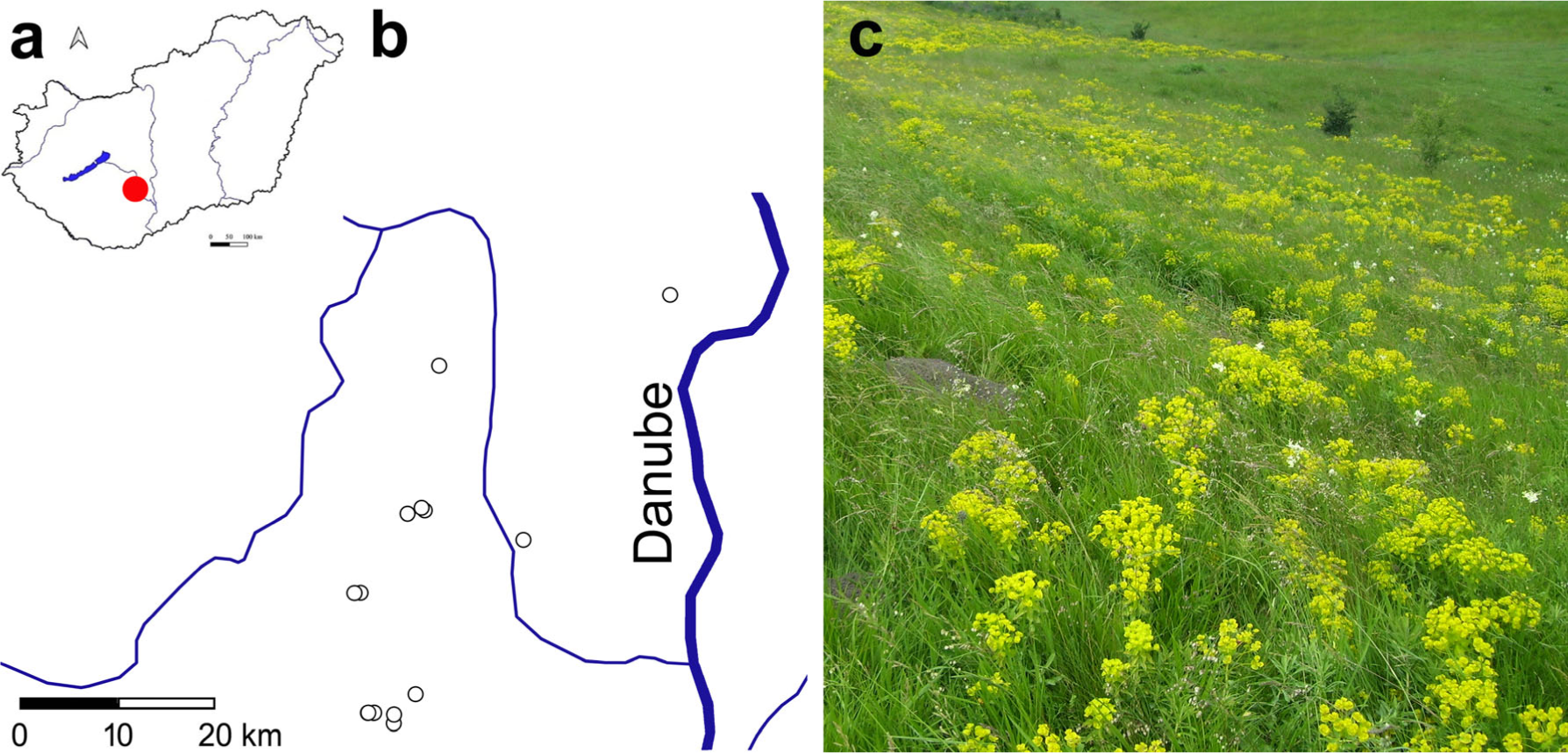
Location of sampling sites (a, b) and an old-growth grassland site (c).

The studied grasslands receive special conservation interest and are included into the Natura 2000 network as a priority habitat type ‘6250 Pannonic loess steppic grasslands’. Loess steppic grasslands are in general indicated by a high richness of forbs and characterised by several tussock forming or stoloniferous grasses (e.g., *Agropyron cristatum*, *Brachypodium pinnatum*, *Bromus erectus* and *B. inermis*, *Chrysopogon gryllus*, *Festuca pseudovina* and *F. rupicola*, and several *Stipa* species, Illyés and Bölöni, 2007). Old-growth grasslands cover various types of loess deposits and were situated in naturally occurring forest-steppe openings (Borhidi et al., 2012; Erdős et al., 2018). The studied loess grasslands in the region cover chernozemic soils, which are developed on loess bedrock. Loess grasslands were historically embedded in a mosaic landscape of the forest-steppe zone (Erdős et al., 2018). The studied landscape was characterised historically by dry loess grasslands, semi-dry grasslands both with and without shrubs, and fragments of open forest-steppes and closed forests (Teleki et al., 2020). Because of the high suitability of chernozemic soils for agricultural production, old-growth loess vegetation has survived on sites inappropriate or difficult for cultivation (e.g., on steep valley sides, road verges or burial mounds, Deák et al., 2016).

In total, 8 old-growth and 5 secondary grassland fragments were selected for a detailed study, ranging from 0.5 to 1 hectare in area. In most of the selected grassland fragments, management ceased decades ago, while some of them have been managed by low intensity or occasional cattle or sheep grazing (Table 1). Old-growth and secondary grasslands were classified using historical maps. Those grasslands were classified as ‘old-growth’, where grassland cover occurred on consecutive maps continuously since the time of the Second Military Survey (1863-64) (Tímár et al., 2006, http://mapire.eu/hu/map/secondsurvey). Secondary grasslands are spontaneously regenerated following the former agricultural use as crop fields, vineyards, or orchards; we selected secondary grasslands older than 50 years. Maps were obtained from the Archives of Tolna county, including maps in manuscript from the end of the 18th to the end of the 19th centuries.

**Table 1.** Detailed information on sampling sites. Origin: O– old-growth grassland, S – secondary grassland.

| Site code | Nearest settlement | Origin | Altitude (m) | Sampling date | GPS coordinates | Area (ha) | Cessation of agricultural cultivation |
| --- | --- | --- | --- | --- | --- | --- | --- |
| FN | Felsőnána | O | 135 | 21.09.2019 | N46.481694 E18.530726 | 0.5 | NA |
| UZ | Uzd | O | 140 | 10.09.2019 | N46.614828 E18.567976 | 1.0 | NA |
| MT | Majos (Temető) | S | 135 | 26.09.2019 | N46.302746 E18.492027 | 0.5 | >50 years |
| MK | Majos (Katonaitemető) | S | 140 | 26.09.2019 | N46.302872 E18.492606 | 0.5 | >50 years |
| TEI | Tevel I | O | 165 | 02.10.2019 | N46.409919 E18.472110 | 0.5 | NA |
| TEII | Tevel II | O | 160 | 03.10.2019 | N46.409605 E18.464368 | 0.5 | NA |
| KTI | Kistormás I | O | 110 | 14.10.2019 | N46.484882 E18.552840 | 1.0 | NA |
| KTII | Kistormás II | O | 115 | 14.10.2019 | N46.487187 E18.548739 | 0.5 | NA |
| BF | BonyhádFáytelep | S | 145 | 16.10.2019 | N46.293873 E18.518098 | 0.5 | >50 years |
| BP | Bonyhád, Prikk-lejtő | S | 135 | 16.10.2019 | N46.301649 E18.518269 | 0.5 | >50 years |
| BL | Bonyhádlőtér | S | 120 | 18.10.2019 | N46.320076 E18.545515 | 1.0 | >50 years |
| SZII | Szedres II | O | 95 | 19.10.2019 | N46.460099 E18.680940 | 1.0 | NA |
| DK | Dunakömlőd | O | 122 | 20.10.2019 | N46.681876 E18.865621 | 1.0 | NA |

### 2.2. Sampling setup

In each grassland site we designated five 10m×10m-sized study areas in which plots of different sizes were embedded. We started the sampling of the sites from one corner and increased the plot size using the embedded plot sampling proposed by Dengler et al. (2016) and Dengler et al. (2018). We sampled the vegetation in plots with the following sizes:1) 10cm×10cm, 2) 30cm×30cm, 3) 50cm×50cm, 4) 1m×1m, 5) 2m×2m, 6) 3m×3m, 7) 4m×4m, 8) 5m×5m, and 9) 10m×10m. In each plot, we recorded the vegetation cover by species on a percentage scale. Nomenclature follows Király et al (2009).

### 2.3. Data capture and analyses

We extracted trait data of species from the Pannonian Database of Plant Traits (PADAPT, Sonkoly et al., 2022). We obtained data for life form, plant height, thousand-seed weight, seed bank type, clonality, rosette formation likeliness, leaf dry weight (LDW), leaf area (LA), specific leaf area (SLA) and leaf dry matter content (LDMC). For further information for the used trait data see Table 2. For clonal spread ability we used the CLO-PLA database and classified the species into four ordinal categories based on potential distance of lateral spread (Klimešová and de Bello 2017, Table 2). We used both classical and functional diversity metrics to compare the vegetation diversity of old growth and secondary grasslands. We calculated species richness (S), Shannon diversity (H, Shannon 1948), Pielou evenness (E, Pielou, 1975) and Berger-Parker dominance index (d, Berger and Parker, 1970). Pielou evenness is calculated with the following equation: E=H/lnS, where E is the Pielou evenness, H is the Shannon diversity, and S is the species richness. The Berger-Parker dominance index is a simple measure of the relative abundance of the most abundant species, and it was calculated with the following equation: d=A_max_/A_total_, where d is the Berger-Parker dominance index, A_max_ is the abundance of the most abundant species, and A_total_ is the summarised abundance of all species present.

**Table 2.** Species traits used in the analyses.

| **Trait** | **Variable type** | **Values** | **Measurement unit** |
| --- | --- | --- | --- |
| Clonal spread | ordinal | 1= no clonal spreading, 2= < 0.01 m/year, 3= 0.01–0.25 m/year, 4= >0.25 m/year. | NA |
| Leaf area (LA) | scale | score | mm<sup>2</sup> |
| Leaf dry matter content (LDMC) | scale | score | mg/g |
| Leaf dry weight (LDW) | scale | score | mg |
| Lifeform | ordinal | 1=annual, 2=biennial, 3=perennial, 4=semi-shrub/chamaephyte, 5= phanerophyte | NA |
| Plant height | scale | score | cm |
| Rosette formation likelihood | binary | 1 = no rosette, 2= rosette (or semi-rosette), | NA |
| Seed bank type | binary | 1=persists seed bank, 0=transient seed bank | NA |
| Specific leaf area (SLA) | scale | score | mm<sup>2</sup>/mg |
| Thousand-seed weight | scale | score | g |

We also compared the variation of diversity indices across spatial scales(scale-dependency)in old-growth and secondary grasslands by studying species area relationships (SAR) using the power function (Arrhenius, 1921; Chiarucci et al., 2006), because it performed well in former comparisons including small plot sizes suggested by several authors (e.g., Turtureanu et al., 2014; Zhang et al., 2021), and its parameters are easy to understand. The power-law function reads as follows: logS = logc + z logA, where S is species richness, A is the sampled area, ‘c’ and ‘z’ are fitted parameters, which represent the number of species per sampled area (c) and the increment of the number of species with the increase of sampling area (z, ‘steepness’ of the curve), respectively. The c and z scores were compared between old-growth and secondary grasslands using a Mann-Whitney U test. The fitting was calculated in R using the ‘sars’ package (R Core Team, 2010; Triantis et al., 2011; Matthews et al., 2022).

We also calculated the three components of functional diversity (functional richness, functional evenness, and functional divergence, Mason et al., 2005; Laliberté and Legendre, 2010). In addition, we calculated functional dispersion (FDis) to measure the functional similarity amongst the characteristic species of the species assemblages; high dispersion scores display high levels of niche differentiation and decreased levels of competition (Mason et al., 2005; Villéger et al., 2008). We calculated the community weighted means (CWMs) of all single traits. All functional diversity calculations were executed cover weighted by using Gower distance measure and the FDiversity programme package (Casanoves et al., 2011). We averaged the scores of each metric at the site level; we averaged scores for plots of the same size originating from the five replications of each site. We analysed the effect of grassland origin (binary variable) and plot size (repeated measure factor, 9 levels) in a two-way repeated measure GLM. We used SPSS 26.0 for the statistical analyses (IBM, 2019). We compared the vegetation composition of the sites using a DCA ordination in CANOCO 5.0 (ter Braak and Smilauer, 2012). Ordination was based on the species abundance data of the largest, 20 m × 20 m sized, plots. Indicator species analysis (IndVal) was applied to identify species typical for old-growth and secondary grasslands (Dufrêne and Legendre, 1997). For the IndVal analysis, we used the ‘labdsv’ package in an R environment (R Core Team, 2010; Roberts, 2010).

## 3. Results

We found that species richness, Shannon diversity and Berger-parker dominance were significantly affected by the grassland origin: while the species richness and Shannon diversity were higher (Table 3., Figure2 and 3), the Berger-Parker dominance was lower in old-growth grasslands (Figure 4). For the multi-trait functional diversity indices, except for functional divergence, no significant effect of grassland origin was found. Analysing the community weighted means of single traits, we found that most traits were significantly affected by grassland origin. While the CWM of lifeform, plant height, thousand-seed weight, leaf dry weight and LDMC was higher, that of seed bank type, rosette formation likeliness and SLA were lower in old-growth grasslands (Figure 4.).

**Table 3.** Effect of grassland origin (old-growth or secondary), plot size and their interaction on the species-, functional diversity, and community weighted mean (CWM) of single traits of the studied grasslands. Significant effects were denoted with **boldface** (*p*<0.05), marginally significant effects (*p*<0.1) with *italics* (Two-way repeated measure GLM, Repeated measure factor was the plot scale).

| Characteristic | Grassland origin |  | Plot size |  | Grassland origin<br>× Plot scale |  |
| --- | --- | --- | --- | --- | --- | --- |
|  | F | *p* | F | *p* | F | *p* |
| **Species diversity** |  |  |  |  |  |  |
| Species richness | **51.144** | **<0.001** | **29.857** | **0.003** | **4.880** | **0.071** |
| Shannon diversity | **63.452** | **<0.001** | **29.892** | **0.003** | 0.865 | 0.602 |
| Berger-Parker dominance | **9.493** | **0.010** | 2.879 | 0.161 | 0.602 | 0.749 |
| Evenness | 1.298 | 0.279 | 2.902 | 0.159 | 1.297 | 0.159 |
| **Functional diversity** |  |  |  |  |  |  |
| Rao entropy | 1.424 | 0.258 | 1.337 | 0.414 | 0.685 | 0.700 |
| Functional richness | 0.626 | 0.445 | 2.317 | 0.217 | 1.729 | 0.313 |
| Functional evenness | 0.666 | 0.432 | **7.018** | **0.039** | *4.347* | *0.086* |
| Functional divergence | **6.953** | **0.023** | 1.082 | 0.505 | **11.471** | **0.016** |
| Functional dispersion | 0.673 | 0.429 | 0.646 | 0.723 | 1.191 | 0.463 |
| **CWM of single traits** |  |  |  |  |  |  |
| Lifeform | **7.599** | **0.019** | 3.863 | 0.104 | 1.166 | 0.104 |
| Plant height | **5.107** | **0.045** | 1.439 | 0.384 | 1.011 | 0.534 |
| Thousand-seed weight | *4.679* | *0.053* | 0.733 | 0.672 | 1.107 | 0.495 |
| Seed bank type | **5.466** | **0.039** | 1.550 | 0.354 | 1.176 | 0.468 |
| Clonality | 1.388 | 0.264 | 1.048 | 0.518 | 1.645 | 0.332 |
| Rosette formation | **5.902** | **0.033** | 2.987 | 0.153 | 1.012 | 0.534 |
| Leaf dry weight (LDW) | *3.909* | *0.074* | 0.569 | 0.770 | 0.486 | 0.821 |
| Leaf area (LA) | 1.208 | 0.295 | 0.706 | 0.688 | 0.671 | 0.708 |
| Specific leaf area (SLA) | **9.314** | **0.011** | 2.069 | 0.252 | 2.269 | 0.223 |
| Leaf dry matter content (LDMC) | **38.435** | **<0.001** | 1.383 | 0.400 | 0.528 | 0.795 |

**Figure 2.**
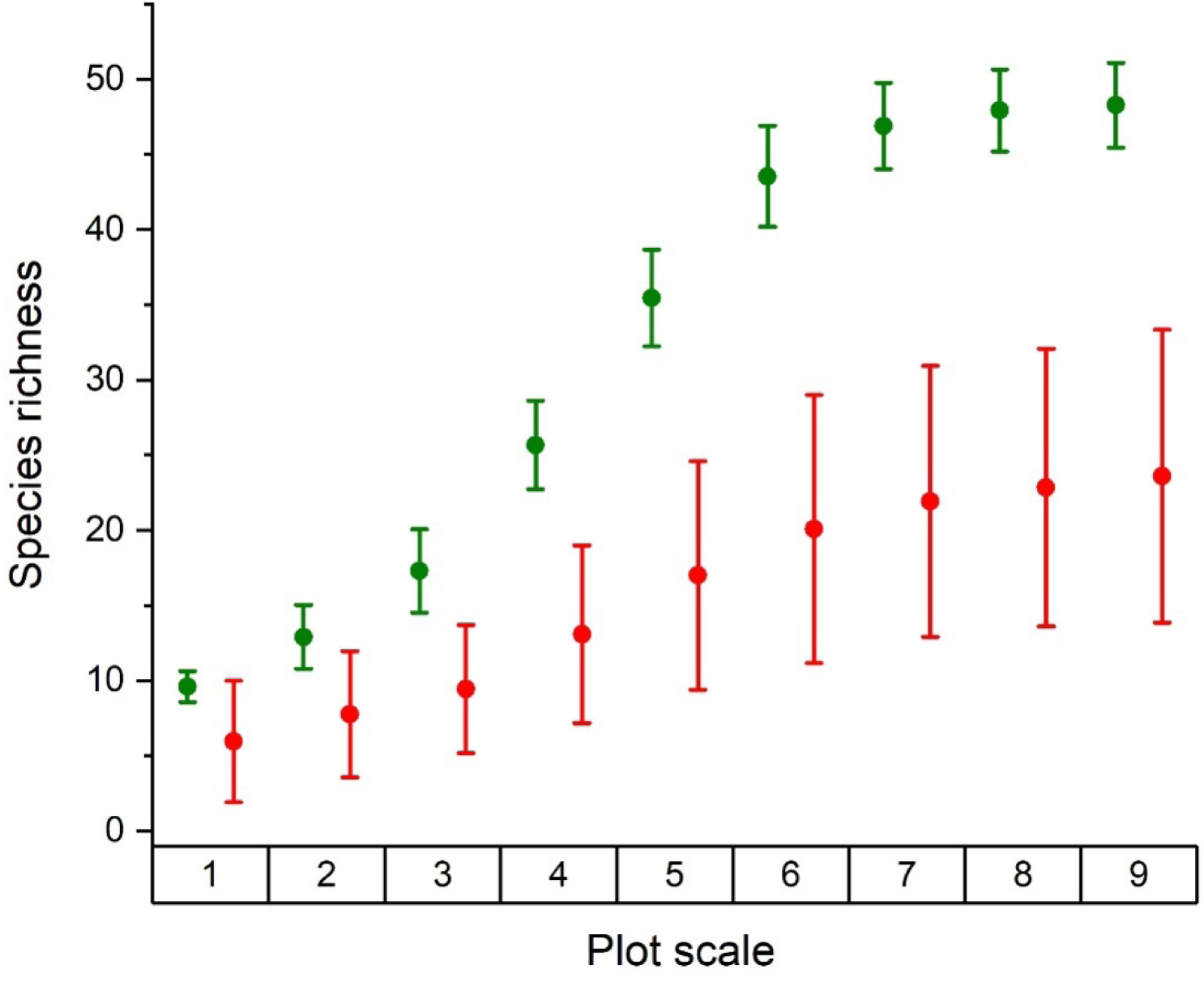
Species richness in old-growth (green) and secondary (red) grasslands (mean±95% CI) across several plot sizes. Species richness is expressed as number of species. The plot sizes were 1) 10×10 cm, 2) 30×30 cm, 3) 50×50 cm, 4) 1×1 m, 5) 2×2 m, 6) 3×3 m, 7) 4×4 m, 8) 5×5 m, and 9) 10×10 m.

**Figure 3.**
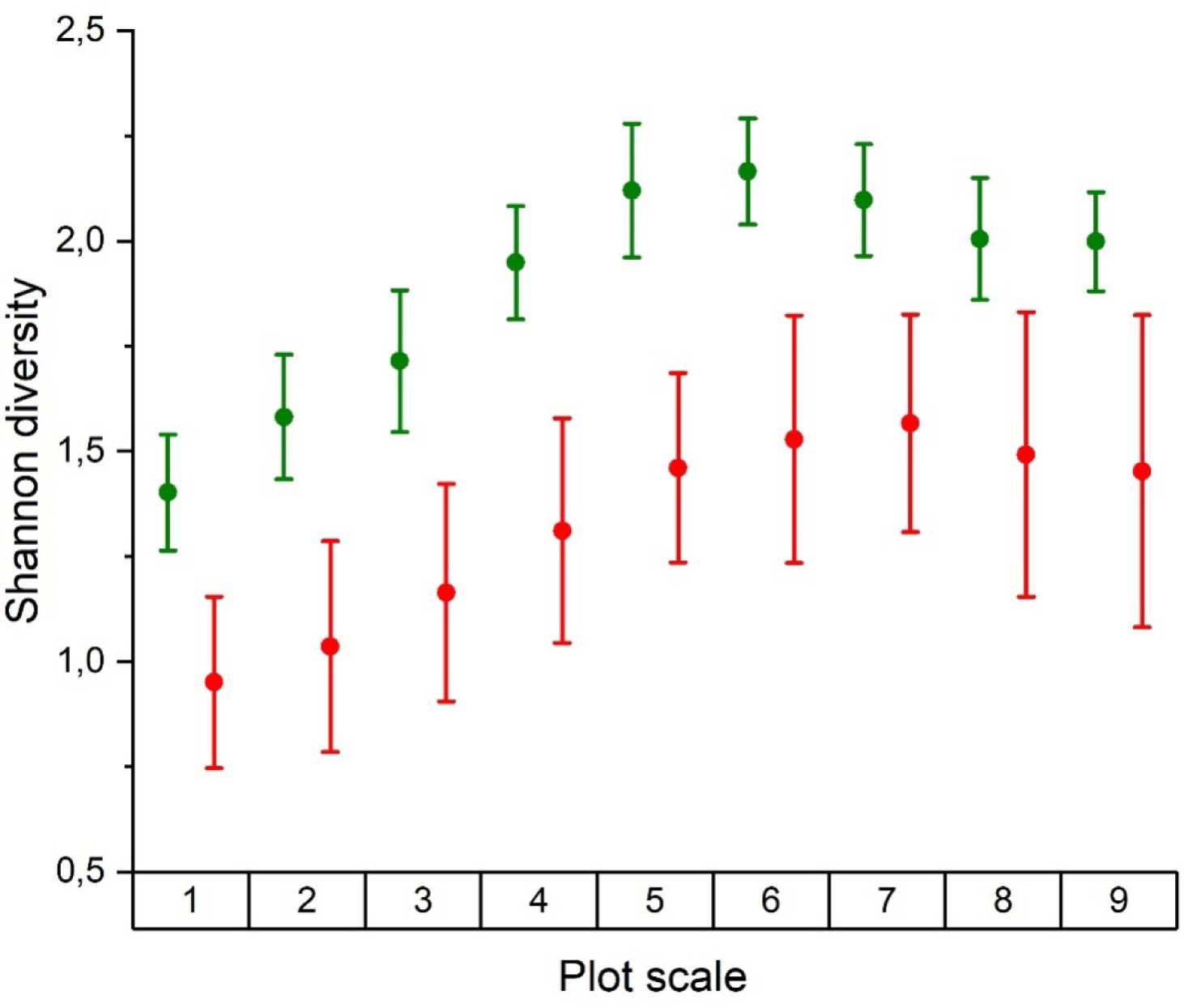
Shannon diversity in old-growth (green) and secondary (red) grasslands (mean±95% CI) across several plot sizes. The plot sizes were 1) 10×10 cm, 2) 30×30 cm, 3) 50×50 cm, 4) 1×1 m, 5) 2×2 m, 6) 3×3 m, 7) 4×4 m, 8) 5×5 m, and 9) 10×10 m.

**Figure 4.**
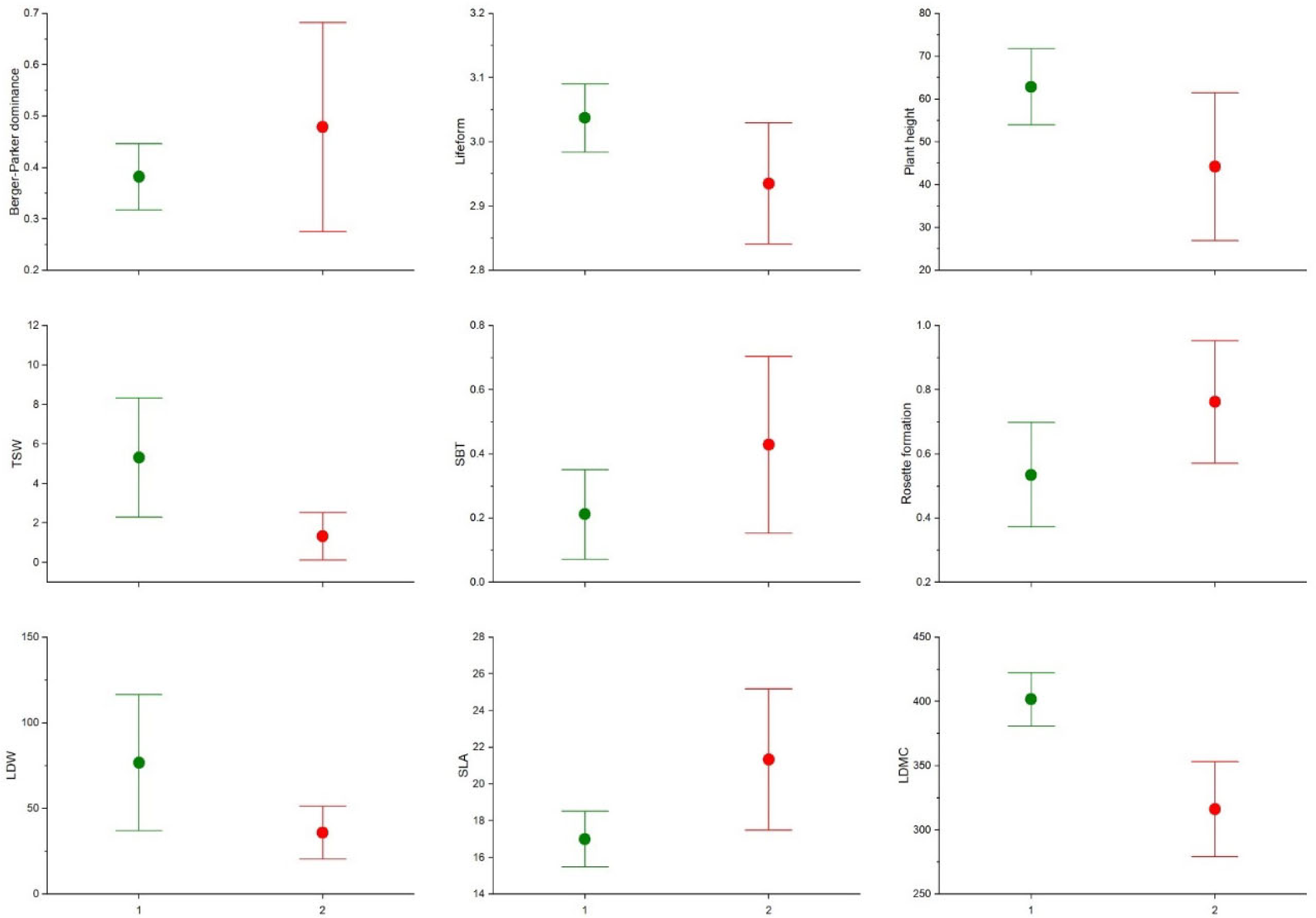
Berger-Parker dominance and community weighted means of various traits in old-growth (green) and secondary (red) grasslands (mean±95% CI). Only those traits were displayed that were proven to be significantly different between old-growth and secondary grasslands in the two-way repeated measure GLM (p<0.05, Table 1).

**Figure 5.**
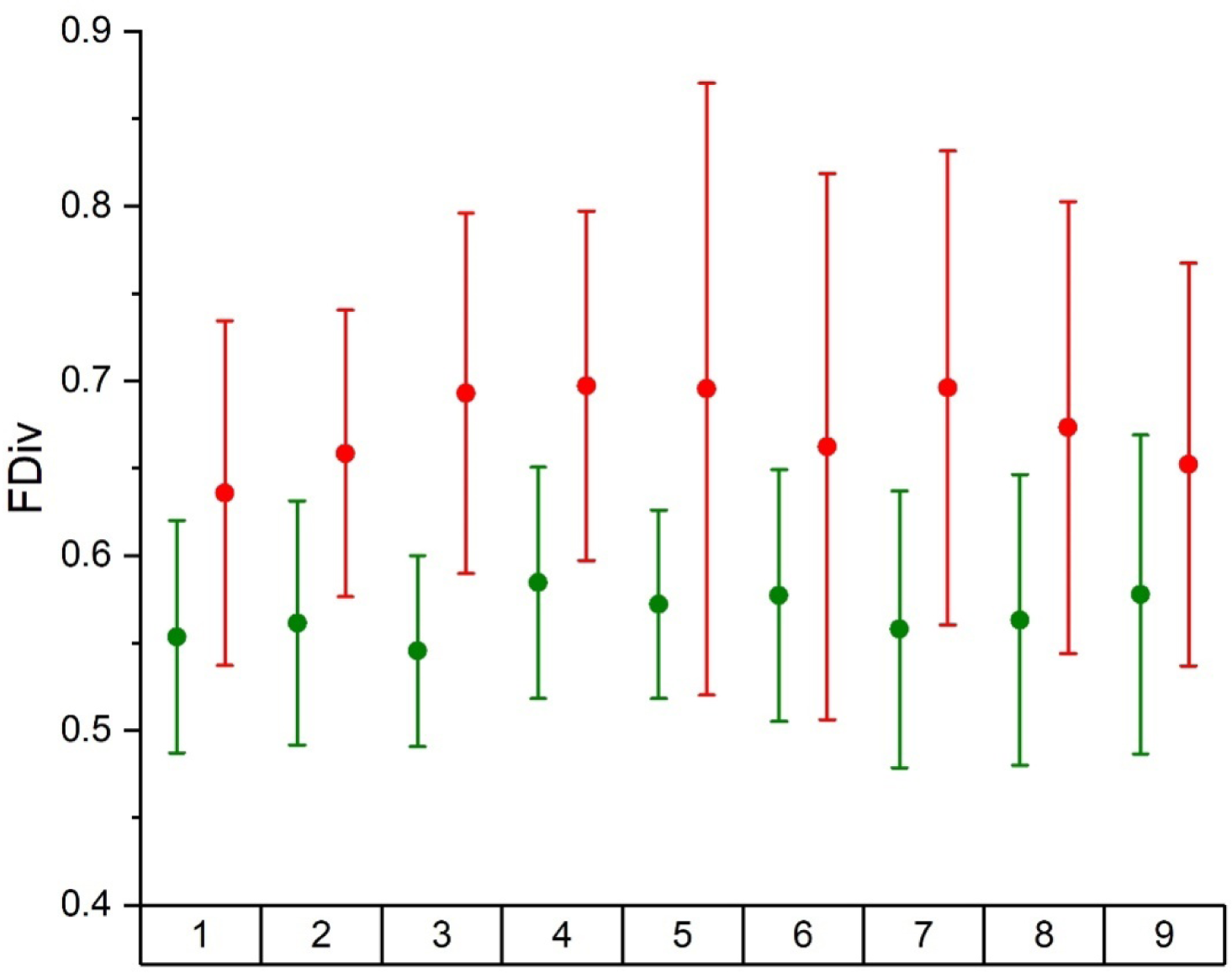
Functional divergence (FDiv) in old-growth (green) and secondary (red) grasslands (mean±95% CI) across several plot sizes. The plot scales were 1) 10×10 cm, 2) 30×30 cm, 3) 50×50 cm, 4) 1×1 m, 5) 2×2 m, 6) 3×3 m, 7) 4×4 m, 8) 5×5 m, and 9) 10×10 m.

Species richness and Shannon diversity displayed significant scale-dependency in the studied grasslands (Table 3). Species richness and Shannon diversity were higher at all plot sizes in old-growth grasslands (Figure 2 and 3). Surprisingly, apart from functional evenness, none of the multi-trait indices or single-trait community weighted means displayed a scale dependency effect (Table 3). We found that the ‘c’ parameter was significantly higher in old-growth grasslands than in secondary ones (*p*<0.001; 3.178 and 2.469, respectively), while the ‘z’ parameter differed only marginally significantly between the grassland types (*p*=0.093, 0.205and 0.174, respectively).

We found that there was a high difference in species composition between old-growth and secondary grasslands. The point clouds of old-growth and secondary grasslands were clearly separated in the DCA ordination (Figure 6). The point clouds of the secondary grasslands were separated along the first, while that of old-growth grasslands along the second DCA axis (Figure 6.). It is clearly indicated also by the DCA ordination that species richness increased from secondary grasslands towards old-growth ones. We identified 52 character species typical for old-growth grasslands and 25 for the secondary ones (Appendix 1). Several species of conservation interest were found among the character species of old-growth grasslands (e.g., *Adonis vernalis*, *Inula germanica*, or *Scabiosa canescens*), while disturbance tolerant species (e.g., *Dactylis glomerata*, *Bellis perennis*, or *Trifolium repens*) or several weeds (e.g., *Taraxacum officinale*, *Setaria viridis*, or *Erodium cicutarium*) were characteristic to secondary grasslands.

**Figure 6.**
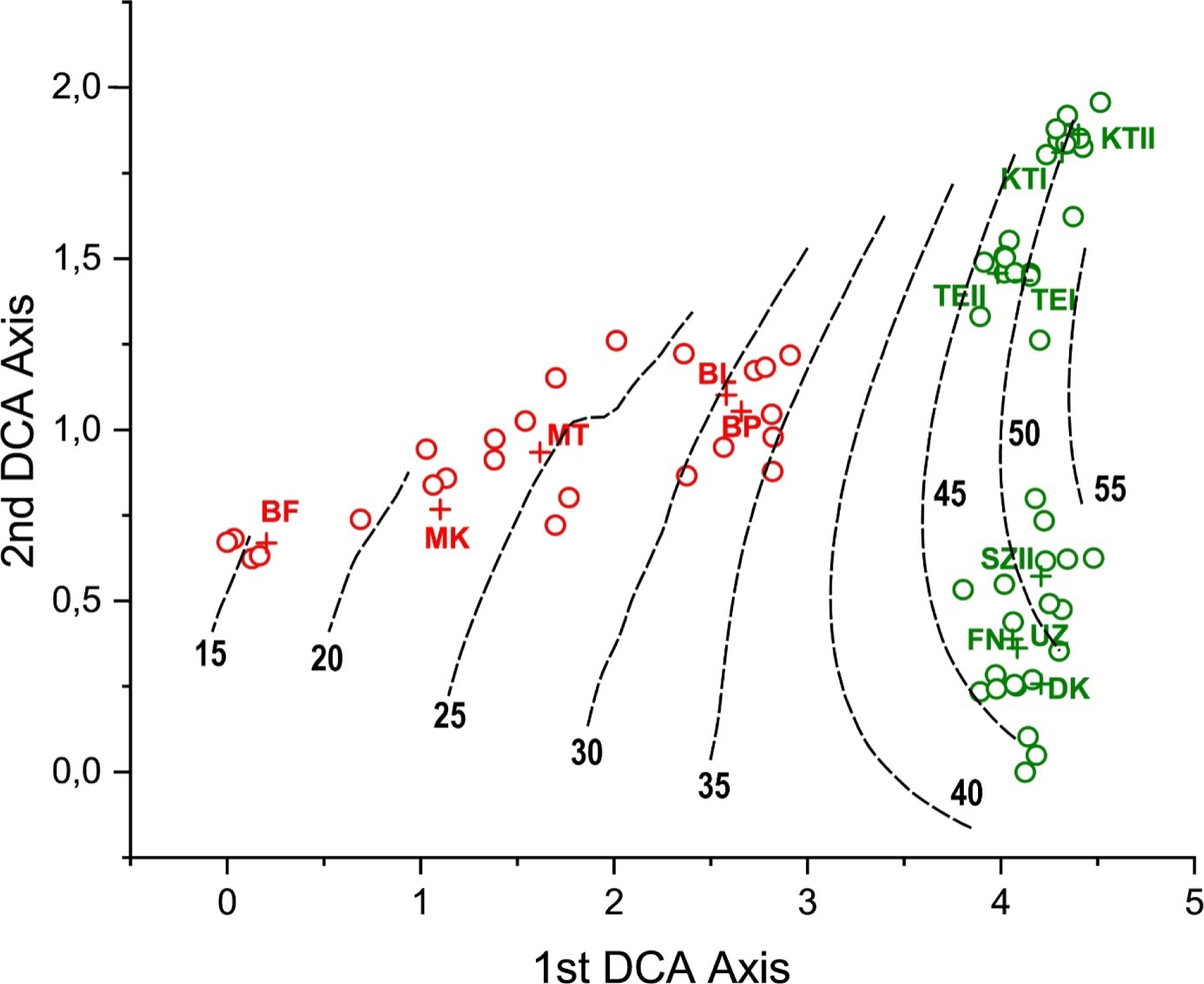
Difference in the species composition of old-growth and secondary grasslands displayed by a DCA ordination (ordination was based on the species abundance data of the largest, 20 m × 20 m sized plots). Eigenvalues of the DCA are 0.696 and 0.245for the first and second axes, respectively. Gradient lengths are 4.51 and 1.96 for the first and second axes, respectively. Cumulative species variance explained by the first four axes is 32.63. Dashed lines represent isolines for species richness (from 15 to 55). Green colours represent old-growth, while red secondary grasslands, with ‘+’ sign centroids of sites are denoted. Site codes are the followings: FN: Felsőnána, UZ: Uzd, MT: Majos (Temető), MK: Majos (Katonaitemető), TEI: Tevel I, TEII: Tevel II, KTI: Kistormás I, KTII: Kistormás II, BF: BonyhádFáytelep, BP: Bonyhád, Prikk-lejtő, BL: Bonyhádlőtér, SZII: Szedres II, DK: Dunakömlőd.

**Appendix 1A.** Indicator species of the old-growth grasslands. Species are identified by an indicator species analysis (IndVal).

| Species | IndVal score | Significance |
| --- | --- | --- |
| *Chrysopogon gryllus* | 1.000 | 0.001 *** |
| *Teucrium chamaedrys* | 0.975 | 0.001 *** |
| *Centaurea scabiosa* | 0.916 | 0.001 *** |
| *Brachypodium pinnatum* | 0.880 | 0.001 *** |
| *Filipendula vulgaris* | 0.880 | 0.001 *** |
| *Medicago falcata* | 0.868 | 0.001 *** |
| *Eryngium campestre* | 0.853 | 0.001 *** |
| *Adonis vernalis* | 0.851 | 0.001 *** |
| *Viola ambigua* | 0.837 | 0.001 *** |
| *Salvia pratensis* | 0.825 | 0.001 *** |
| *Crataegus monogyna* | 0.822 | 0.001 *** |
| *Asperula cynanchica* | 0.814 | 0.001 *** |
| *Knautia arvensis* | 0.759 | 0.002 ** |
| *Pseudolysimachion spicatum* | 0.758 | 0.002 ** |
| *Taraxacum serotinum* | 0.742 | 0.001 *** |
| *Nonea pulla* | 0.725 | 0.001 *** |
| *Botriochloa ischaemum* | 0.710 | 0.013 * |
| *Hypericum perforatum* | 0.707 | 0.005 ** |
| *Galium glaucum* | 0.707 | 0.001 *** |
| *Euphorbia glareosa* | 0.689 | 0.001 *** |
| *Tanacetum corymbosum* | 0.689 | 0.004 ** |
| *Sanguisorba minor* | 0.685 | 0.002 ** |
| *Seseli anuum* | 0.662 | 0.001 *** |
| *Cirsium boujartii* | 0.652 | 0.002 ** |
| *Trifolium montanum* | 0.652 | 0.001 *** |
| *Campanula bononiensis* | 0.632 | 0.001 *** |
| *Thymus glabrescens* | 0.632 | 0.007 ** |
| *Inula germanica* | 0.612 | 0.001 *** |
| *Serratula tinctoria* | 0.612 | 0.002 ** |
| *Anchusa officinalis* | 0.592 | 0.005 ** |
| *Ononis spinosa* | 0.592 | 0.002 ** |
| *Chamaenerion angustifolium* | 0.570 | 0.004 ** |
| *Elymus hispidus* | 0.569 | 0.039 * |
| *Lathyrus tuberosus* | 0.556 | 0.015 * |
| *Stipa capillata* | 0.548 | 0.008 ** |
| *Astragalus onobrychis* | 0.548 | 0.007 ** |
| *Peucedanum alsaticum* | 0.548 | 0.006 ** |
| *Thalictrum minus* | 0.524 | 0.007 ** |
| *Rapistrum perenne* | 0.524 | 0.011 * |
| *Briza media* | 0.524 | 0.005 ** |
| *Potentilla erecta* | 0.514 | 0.031 * |
| *Scabiosa canescens* | 0.500 | 0.025 * |
| *Artemisia campestris* | 0.500 | 0.010 ** |
| *Betonica officinalis* | 0.500 | 0.026 * |
| *Rosa canina* | 0.500 | 0.010 ** |
| *Senecio jacobea* | 0.500 | 0.008 ** |
| *Stachys recta* | 0.500 | 0.010 ** |
| *Verbena officinalis* | 0.474 | 0.022 * |
| *Senecio erucifolius* | 0.447 | 0.039 * |
| *Linum austriacum* | 0.447 | 0.016 * |
| *Prunella laciniata* | 0.447 | 0.013 * |
| *Allium sphaerocephalon* | 0.418 | 0.037 * |

**Appendix 1B.** Indicator species of the secondary grasslands. Species are identified by an indicator species analysis (IndVal).

| Species | IndVal score | Significance |
| --- | --- | --- |
| *Achillea collina* | 0.963 | 0.001 *** |
| *Taraxacum officinale* | 0.883 | 0.001 *** |
| *Cynodon dactylon* | 0.872 | 0.001 *** |
| *Trifolium pratense* | 0.807 | 0.001 *** |
| *Dactylis glomerata* | 0.791 | 0.002 ** |
| *Plantago lanceolata* | 0.771 | 0.012 * |
| *Glechoma hederacea* | 0.721 | 0.001 *** |
| *Leontodon hispidus* | 0.660 | 0.031 * |
| *Astragalus cicer* | 0.650 | 0.008 ** |
| *Clinopodium vulgare* | 0.646 | 0.024 * |
| *Viola odorata* | 0.632 | 0.001 *** |
| *Bellis perennis* | 0.600 | 0.001 *** |
| *Galium mollugo* | 0.569 | 0.005 ** |
| *Setaria viridis* | 0.566 | 0.002 ** |
| *Brachypodium sylvaticum* | 0.565 | 0.001 *** |
| *Potentilla arenaria* | 0.560 | 0.034 * |
| *Erodium cicutarium* | 0.529 | 0.001 *** |
| *Erigeron annuus* | 0.517 | 0.005 ** |
| *Cichorium intybus* | 0.447 | 0.004 ** |
| *Oxalis corniculata* | 0.447 | 0.009 ** |
| *Prunus spinosa* | 0.447 | 0.007 ** |
| *Trifolium repens* | 0.447 | 0.009 ** |
| *Calamagrostis epigeios* | 0.447 | 0.007 ** |
| *Tragopogon orientalis* | 0.432 | 0.025 * |
| *Potentilla reptans* | 0.346 | 0.045 * |

## 4. Discussion

### 4.1. Unsuccessful recovery of species diversity in secondary grasslands

We found that species richness and Shannon diversity were higher in old-growth grasslands, emphasizing that species diversity cannot recover within 50 years after the destruction of primeval grasslands. Similar results have been found by some other studies dealing with spontaneous secondary succession in loess and other types of steppic grasslands (e.g., Molnár and Botta-Dukát, 1998). One of the potential causes may be the very slow immigration rate of specialist species in old fields. Molnár and Botta-Dukát (1998) revealed that some of the grassland specialists, like *Potentilla arenaria*, *Filipendula vulgaris*, *Fragaria viridis* or *Thalictrum minus*, couldn’t been detected in several-decades-old fields, despite of their high abundance in the surrounding landscape (Molnár and Botta-Dukát, 1998). In spontaneously recovering alfalfa fields, generalist species colonised quite rapidly (within 10 years) after abandonment, while several grassland specialists, including characteristic forbs, were still missing from the secondary grasslands and the species richness was lower than in reference old-growth grasslands (Török et al., 2011b). Other research by Ruprecht (2006) found contrasting results, and provided information on very successful establishment of dry grassland species in Romanian old-fields. It was found that in landscapes with a high proportion of species-rich grasslands thus abundant propagule sources, only a very small fraction of target species was missing from the vegetation of the secondary grasslands after a couple of decades.

Another cause of the unsuccessful recovery of species diversity may be the increasing dominance by generalist species with a high competitive ability, especially grasses, in the mid-phase of secondary succession (i.e., a few decades after the initiation of secondary succession), which “closes” the colonisation window and slows down or even blocks further recovery (Bartha et al., 2003; Albert et al., 2014; Bartha et al., 2014). This is in accordance with our finding regarding the Berger-Parker dominance, which was higher in secondary compared to old-growth grasslands, suggesting high competitive pressure by one very abundant species in secondary grasslands.

### 4.2. Secondary grasslands are functionally as diverse as old-growth grasslands

Based on similar multi-trait functional diversity indices of secondary and old-growth grassland stands, we infer that dry grasslands in our study area could reassemble relatively quickly (within 50 years) to well-functioning systems. Tölgyesi et al. (2019) found similar patterns in spontaneously secondary sand and loess grasslands and proposed that functional diversity may recover much faster than species diversity. Rapidly colonising species may fill up important functions (niches) in reassembled grassland communities; however, functional redundancy remains to be waited for even after several decades. The slow immigration rate of species, thus low species richness even after 50 years may be responsible for the lower functional divergence thus low functional redundancy of secondary grasslands. In the face of future climate change or disturbances, functional diversity and functional redundancy of grassland plant communities are of outstanding importance. The disappearance of certain species will be less harmful to the functioning of the system if other species with similar traits take over their role.

However, our results only partly confirmed the first hypothesis, as for the multi-trait functional diversity indices, except for functional divergence, no significant effect of grassland origin was detected. Studying sand grassland and loess steppe restorations, Tölgyesi et al. (2019) found that while functional diversity of spontaneously secondary sites became very similar to that of reference old-growth grasslands, species richness was significantly higher in the latter. These results are in line with our findings as we found the same for multi-trait indices except for functional divergence. One likely explanation of the absence of significant differences between functional richness and evenness in old-growth and restored grasslands is provided also by Tölgyesi et al. (2019). The study stressed that it is likely that functional diversity recovers much faster than species diversity. In contrast to the results of Tölgyesi et al. (2019) we found that the functional divergence in secondary grasslands was higher compared to old-growth grasslands and also its variance was much higher. It was found that both the success in the recovery of a characteristic species assembly is highly site dependent (Ruprecht, 2006), and also the formation of a mid-succession grass-dominance is highly species- and site-specific (Bartha et al., 2014). High functional divergence means, that the most abundant species are characterised with extremities of the functional character range and the niche differentiation is high (Mason et al., 2005). A likely explanation for high functional divergence and its high variance in secondary grasslands can be explained by lower stability of secondary grasslands compared to that of old-growth grasslands; it was found in former research that disturbance-driven immigration and disappearance of species are likely more common in secondary grasslands than in old-growth grasslands (Bartha et al., 2003; Virágh and Bartha, 2003).

### 4.3. Scale dependency of diversity patterns

We found that opposite to functional diversity metrics except functional evenness, species richness and Shannon diversity were highly scale-dependent. However, old-growth grasslands were more diverse than secondary grasslands across all the studied plot sizes. This finding was strengthened by the higher ‘c’ parameter of the power function, representing species-area relationships, in old-growth grasslands. These results do not support our second hypothesis about the scale-dependency of differences in taxonomic and functional diversity measures between the two grassland types. The ‘z’ parameter of the power function representing species-area relationships was marginally significantly higher in the case of old-growth stands, showing that the increment in species number with increasing plot size is faster in old-growth grasslands. Thus, larger-scale taxonomic diversity is disproportionally higher in old-growth compared to secondary grasslands, which means that old-growth grasslands are more heterogeneous in respect of their species composition, thus probably have higher beta-diversity compared to secondary stands (Crist and Veech, 2006). For ‘z’ values, our findings (0.174-0.205) fit well into the range reported in a loess grassland study with similar plot sizes by Turtureanu et al. (2014) (0.159-0.264). Similar ‘z’ values were obtained in other studies of dry grasslands and mesic meadows (Dengler, 2005; Chiarucci et al., 2006; Dolnik and Breuer, 2008). However, much higher ‘z’ values were reported in a study of loess grassland patches on kurgans by Dembicz et al. (2020) (0.32), although the area range was different (between 107 and 4449 m^2^).

### 4.4. Compositional differences between old-growth and secondary grasslands

We hypothesised that there is a high compositional difference between primary and secondary grasslands. This hypothesis was confirmed by our results. It was also found by Nerlekar & Veldman (2020) that the species assembly process in recovering the habitat specific species pool of old-growth grasslands is slow, and high persistence of weedy species is typical in secondary grasslands. In the present study, we found much higher number of characteristic species for old-growth grasslands than for secondary ones, and most characteristic species of secondary grasslands were disturbance tolerant species or weeds. This pattern was also found by former research analysing successional patterns in secondary steppic loess and sand grasslands (Török et al., 2011b; Bartha et al., 2014; Albert et al., 2014).

However, not only was the species composition highly distinct, but also most of the community weighted means of single traits were significantly different between old-growth and secondary grasslands. We found that old-growth grasslands are characterised by taller, larger seeded, perennial species, having higher leaf dry weight and leaf dry matter content. By contrast, secondary grasslands are characterized by rosette-forming species, having high SLA and forming persistent seed banks. In line with these, Veldman et al. (2015) highlighted that old-growth grasslands are characterised by species having a long lifespan, low success of establishing from seeds, forming transient seed banks but persistent underground bud banks.

In dynamically stable communities, long life span, intensive vegetative growth and reproduction, and other characteristics granting competitive advantage over others, like large plant height and large leaf area, will be favoured (Janečková et al., 2017; Deák et al., 2021). These were also supported by the present study. Higher leaf dry matter content and lower specific leaf area are adaptations to drought tolerance in harsh environments (Deák et al., 2021; Lindborg et al., 2014). We found that LDMC was higher while SLA was lower in old-growth grasslands. Rosette forming leaves are close to the soil surface and can capture light effectively when the community biomass is low. Higher likeliness of rosette formation is beneficial in highly disturbed habitats, where biomass production is rather low, or the vegetation is intensively removed and trampled by mowing or intensive livestock grazing (Tóth et al., 2018).

Among the important drivers of compositional differences between old-growth and secondary grasslands we can mention the differences in seed mass and seed bank strategies. We found that in old-growth grasslands, the community weighted mean of seed persistence index was much lower, while that of seed mass was significantly higher than in recovering ones. Weeds and disturbance tolerant species likely build up persistent seed banks, while characteristic perennial and stress tolerant species of pristine grasslands generally have transient seed banks. This was validated for many grassland types of Europe, including also dry grasslands and mesophilous meadows (Bossuyt and Honnay, 2008; Kiss et al., 2016). Under more stable conditions, when the gap formation by disturbance is less frequent, the production of large but rather transient seeds is more likely favoured over the small, easily dispersed, and persistent ones (Bossuyt and Honnay, 2008; Lindborg et al., 2014). Larger seeds grant competitive advantage during germination; higher reserves in the large seeds enable a faster seedling development and larger seedlings, which provides advantage in early establishment and seedling competition (Leishman et al., 2000). Larger seeds can also germinate either from deeper soil layers, or, because of the higher seed reserves, seedlings can breach a thicker litter layer than smaller ones, while reserves enable higher drought tolerance for the seedlings (Janečková et al., 2017; Sonkoly et al., 2020).

## 5. Conclusions

Our results indicate that there are remarkable differences in taxonomic and functional diversity between old-growth grasslands and secondary ones. These differences are clear even after 50 years of secondary succession, indicating that the recovery of species rich loess grasslands requires long time. Our results also point out that during secondary succession there is a likely functional saturation of the species assembly in the first few decades of recovery, and while patterns and structure of secondary grasslands became quite similar to those of old-growth grasslands, the species richness and diversity remains still much lower. Old-growth grasslands support considerable plant diversity, and species composition is slow to recover if destroyed for agricultural land use. This underlines the priority of protecting existing old-growth grassland remnants over restoration actions and the recovery of secondary grasslands. Thus, restoration actions should focus first to enlarge the area of old-growth grassland fragments, to buffer them with secondary grasslands mitigating the negative influence of the surrounding croplands and other human made habitats and embed them into a less hostile matrix. It was also an important finding that, while some taxonomic diversity metrics like Shannon diversity or species richness displayed a high scale dependency, functional characteristics including single trait CWMs did not. This means that functional diversity metrics provide a good basis for comparison of grasslands sampled with different plot sizes.

### Funding sources

PT was supported by the National Research, Development and Innovation Office [KKP144068; K137573] during the manuscript preparation. The field research was financed by the Higher Education Institutional Excellence Programme of the Ministry for Innovation and Technology in Hungary, within the framework of the 3. thematic programme of the University of Pécs. This project has received funding from the HUN-REN Hungarian Research Network.

### CRediT authorship contribution statement

**Péter Török**: Project administration, Funding acquisition, Conceptualization, Formal analysis, Investigation, Methodology, Writing – original draft, Writing – review & editing. **Balázs Teleki**: Investigation, Methodology, Conceptualization, Writing – original draft, Writing – review & editing. **László Erdős**: Methodology, Writing – review & editing. **McIntosh-BudayAndrea**: Investigation, Formal analysis, Writing – review & editing. **Eszter Ruprecht**: Methodology, Writing – original draft, Writing – review & editing. **Béla Tóthmérész**: Conceptualization, Formal analysis, Supervision, Funding acquisition, Methodology, Writing – review & editing.

### Data availability

Data will be made available on request.

## Acknowledgements

Authors are thankful to C. Tölgyesi, Z. Molnár, Z. Bátori for their advises and help.

## References

Albert, Á.-J., Kelemen, A., Valkó, O., Miglécz, T., Csecserits, A., Rédei, T., Deák, B., Tóthmérész, B., Török, P., 2014. Secondary succession in sandy old-fields: a promising example of spontaneous grassland recovery. Appl. Veg. Sci. 17, 214–224. 10.1111/avsc.12068

Arrhenius, O., 1921. Species and area. J. Ecol. 9, 95–99.

Bartha, S., Meiners, S.J., Pickett, S.T.A., Cadenasso, M.L., 2003. Plant colonizationwindows in a mesic old field succession. Appl. Veg. Sci. 6, 205–212.10.1111/j.1654-109X.2003.tb00581.x

Bartha, S., Szentes, S., Horváth, A., Házi, J., Zimmermann, Z., Molnár, C., Dancza, I., Margóczi, K., Pál, R.W., Purger, D., Schmidt, D., Óvári, M., Komoly, C., Sutyinszki, Z., Szabó, G., Csathó, A.I., Juhász, M., Penksza, K. and Molnár, Z. (2014), Impact of mid-successional dominant species on the diversity and progress of succession in regenerating temperate grasslands. Appl. Veg. Sci. 17, 201–213. 10.1111/avsc.12066

Berger, W.H., Parker, F. L., 1970. Diversity of Planktonic Foraminifera in Deep Sea Sediments. Science. 168, 1345–1347. 10.1126/science.168.3937.1345.

Borhidi, A., Kevey, B., Lendvai, G., 2012. Plant communities of Hungary. Budapest, Hungary: Akadémiai Kiadó.

Bossuyt, B., Honnay, O., 2008. Can the seed bank be used for ecological restoration? An overview of seed bank characteristics in European communities. J. Veg. Sci. 19, 875–884. 10.3170/2008-8-18462

ter Braak, C. J. F., Šmilauer, P., 2012. Canoco reference manual and user’s guide: software for ordination, version 5.0. Microcomputer Power.

Buisson, E., Archibald, S., Fidelis, A., Suding, K.N., 2022. Ancient grasslands guide ambitious goals ingrassland restoration. Science. 377, 594–598. 10.1126/science.abo4605

Casanoves, F., Pla, L., Di Rienzo, J.A., Díaz, S., 2011. FDiversity: a software package for the integrated analysis of functional diversity. Methods Ecol. Evol. 2, 233–237. 10.1111/j.2041-210X.2010.00082.x

Cousins, S.A.O., Ohlson, H., Eriksson, O., 2007. Effects of historical and presentfragmentation on plant species diversity in semi-natural grasslands in swedish rurallandscapes. Landsc. Ecol. 22, 723–730. 10.1007/s10980-006-9067-1

Chiarucci, A., Viciani, D., Winter, C., Diekmann, M., 2006. Effects of productivity on speciesarea curves in herbaceousvegetation: evidence from experimental and observational data. Oikos. 115, 475–483. 10.1111/j.2006.0030-1299.15116.x

Crist, T.O., Veech, J.A., 2006. Additive partitioning of rarefaction curves and species–area relationships: unifying α-, β-and γ-diversity with sample size and habitat area. Ecol. Lett. 9, 923–932. 10.1111/j.1461-0248.2006.00941.x

Deák, B., Tóthmérész, B., Valkó, O., Sudnik-Wójcikowska, B., Moysieyenko, I.I., Bragina, T.M., Apostolova, I., Dembicz, I., Bykov, N.I., Török, P., 2016. Cultural monuments and nature conservation: A review of the role of kurgans in the conservation and restoration of steppe vegetation. Biodivers. Conserv. 25, 2473–2490. 10.1007/s10531-016-1081-2

Deák, B., Á. Bede, Z. Rádai, B. Tóthmérész, P. Török, D. Nagy D, A. Torma, G. Lőrinczi, A. Nagy, S. Mizser, A. Kelemen, and O. Valkó. 2021. Different extinction debts among plants and arthropods after loss of grassland amount and connectivity. Biol. Conserv. 264, 109372. 10.1016/j.biocon.2021.109372

Deák, B., Rádai, Z., Lukács, K., Kelemen, A., Kiss, R., Bátori, Z., Kiss, P.J., Valkó, O., 2020. Fragmented dry grasslands preserve unique components of plant species and phylogenetic diversity in agricultural landscapes. Biodivers. Conserv. 29, 4091–4110. 10.1007/s10531-020-02066-7

Deák, B., Rádai, Z., Bátori, Z., Kelemen, A., Lukács, K., Kiss, R., Maák, I.E., Valkó, O., 2021. Ancient Burial Mounds Provide Safe Havens for Grassland Specialist Plants in Transformed Landscapes—A Trait-Based Analysis. Front. Ecol. Evol. 9, 619812. 10.3389/fevo.2021.619812

Dembicz, I., Moysiyenko, I.I., Kozub, Ł., Dengler, J., Zakharova, M., Sudnik-Wójcikowska, B., 2020. Steppe islands in a sea of fields: Where island biogeography meets the reality of a severely transformed landscape. J. Veg. Sci. 32, e12930. 10.1111/jvs.12930

Dengler, J., 2005. Zwischen Estland und Portugal—Gemeinsamkeiten und Unterschiede der Phytodiversitätsmuster Europäischer Trockenrasen. Tuexenia 25, 387–405.

Dengler, J., Janišová, M., Török, P., Wellstein, C., 2014. Biodiversity of palaearctic grasslands: a synthesis. Agric. Ecosyst. Environ. 182, 1–14. 10.1016/j.agee.2013.12.015

Dengler, J., Boch, S., Filibeck, G., Chiarucci, A., Dembicz, I., Guarino, R., Henneberg, B., Janišová, M., Marcenò, C., Naqinezhad, A., Polchaninova, N.Y., Vassilev, K., Biurrun, I., 2016. Assessing plant diversity and composition in grasslands across spatial scales: the standardised EDGG sampling methodology. Bulletin of the Eurasian Dry Grassland Group 32, 13–30.

Dengler, J., Biurrun, I., Dembicz, I., 2018. Standardised EDGG methodology for sampling grassland diversity: second amendment. Palaearctic Grasslands. 49, 22–26. 10.21570/EDGG.PG.49.22-26

Dolnik, C., Breuer, M., 2008. Scale Dependency in the Species-Area Relationship of Plant Communities. Folia Geobot. 43, 305–318. 10.1007/s12224-008-9019-4

Dufrêne, M., Legendre, P., 1997. Species assemblages and indicator species: the need for a flexible asymmetrical approach. Ecol. Monogr. 67, 345–366. 10.1890/0012-9615(1997)067[0345:SAAIST]2.0.CO;2

Erdős, L., Ambarlı, D., Anenkhonov, O.A., Bátori, Z., Cserhalmi, D., Kiss, M., Kröel-Dulay, G., Liu, H., Magnes, M., Molnár, Z., Naqinezhad, A., Semenishchenkov, Y.A., Tölgyesi, C., Török, P., 2018. The edge of two worlds: Eurasian forest-steppes in dynamic transition. Appl. Veg. Sci. 21, 345–362. 10.1111/avsc.12382

European Union, 1992. Council Directive 92/43/EEC of 21 May 1992 on the conservation of natural habitats and of wild fauna and flora.

Hewitt, J. E., Thrush, S. F., Lundquist, C., 2017. Scale-dependence in Ecological Systems. In: eLS. John Wiley & Sons, Ltd: Chichester. 10.1002/9780470015902.a0021903.pub2

IBM, 2019. SPSS Statistics for Windows, Version 26.0. Armonk, NY: IBM Corp.

Illyés, E., Bölöni, J. 2007. Slope steppes, loess steppes and forest steppe meadows in Hungary. Vácrátót, Hungary: MTA Ökológiai és Botanikai Kutatóintézet.

Janečková, P., Janeček, Š., Klimešová, J., Götzenberger, L., Horník, J., Lepš, J., de Bello, F., 2017. The plant functional traits that explain species occurrence across fragmented grasslands differ according to patch management, isolation, and wetness. Landscape Ecol. 32, 791–805. 10.1007/s10980-017-0486-y

Király, G. (Eds.), 2009. New Hungarian Herbal. The Vascular Plants of Hungary (in Hungarian). Identification keys. Jósvafő: Aggtelek National Park Directorate.

Kiss, R., Valkó, O., Tóthmérész, B., Török, P., 2016. Seed bank research in Central-European grasslands - An overview. In: Seed Banks: Types, Roles and Research. Nova Science Publisher, pp. 1-34. ISBN 978-1-536 10-388-5

Klaus, V.H., 2013. Urban Grassland Restoration: A Neglected Opportunity for Biodiversity Conservation. Restor. Ecol. 21, 665–669. 10.1111/rec.12051

Klimešová, J., Danihelka, J., Chrtek, J., de Bello, F., Herben, T., 2017. CLO-PLA: a database of clonal and bud-bank traits of the Central European flora. Ecology 98, 1179. 10.1002/ecy.1745

Laliberté, E., Legendre, P., 2010. A distance-based framework for measuring functional diversity from multiple traits. Ecology. 91, 299–305. 10.1890/08-2244.1

Leishman, M.R. Wright, I.J., Moles, A.T., Westoby, M., 2000. The evolutionary ecology of seed size, in: Fenner M (Eds.), Seeds – the ecology of regeneration in plant communities, vol 2. CAB International, Wallingford, pp 31–57.

Lindborg, R., Plue, J., Andersson, K., Cousins, S.A.O., 2014. Function of small habitat elements for enhancing plant diversity in different agricultural landscapes. Biol. Conserv. 169, 206–213. 10.1016/j.biocon.2013.11.015

Lindgren, J.P., Cousins, S.A.O., 2017. Island biogeography theory outweighs habitat amount hypothesis in predicting plant species richness in small grassland remnants. Landscape Ecol. 32, 1895–1906. 10.1007/s10980-017-0544-5

Löffler, F., Poniatowski, D., Fartmann, T., 2020. Extinction debt across three taxa in well connected calcareous grasslands. Biol. Conserv. 246, 108588. 10.1016/j.biocon.2020.108588

Mason, N.W., Mouillot, D., Lee, W.G., Wilson, J.B., 2005. Functional richness, functional evenness and functional divergence: The primary components of functional diversity. Oikos. 111, 112–118. 10.1111/j.0030-1299.2005.13886.x

Matthews, T.J., Guilhaumon, F., Cazelles, K., 2022. The ‘sars’ R Package. Fit and compare species-area relationship models using Multimodel Inference, retrieved from https://txm676.github.io/sars/

MacArthur, R.H., Wilson, E.O., 1967. The Theory of Island Biogeography. Princeton, NJ: Princeton University Press.

Mezősi, G., 2017. The physical geography of Hungary, 1st ed. Berlin, Germany: Springer.

Molnár, Z., Botta-Dukát, Z., 1998. Improved space-for-time substitution for hypothesis generation: secondary grasslands with documented site history in SE-Hungary. Phytocoenologia. 28, 1–29.

Nerlekar, A.N., Veldman J.W., 2020. High plant diversity and slow assembly of old-growth grasslands. Proc. Nat. Acad. Sci. U S A. 117, 18550–18556. 10.1073/pnas.1922266117

Ottaviani, G., Keppel, G., Götzenberger, L., Harrison, S., Opedal, Ø.H., Conti, L., Liancourt, P., Klimešová, J., Silveira, F.A.O., Jiménez-Alfaro, B., Negoita, L., Doležal, J., Hájek, M., Ibanez, T., Méndez-Castro, F.E., Chytrý, M., 2020. Linking Plant Functional Ecology to Island Biogeography. Trends Plant Sci. 25, 329–339. 10.1016/j.tplants.2019.12.022

Palmer, M.W., White, P.S., 1994. Scale dependence and the species-area relationship. Am. Nat. 144, 717–740.

Pielou, E.C., 1975. Ecological Diversity. New York: Wiley.

Prach, K., Fajmon, K., Jongepierová, I., Řehounková, K. 2015. Landscape context in colonization of restored dry grasslands by target species. Appl. Veg. Sci. 18, 181–189. 10.1111/avsc.12140

Prach, K., Tichý, L., Lencová, K., Adámek, M., Koutecký, T., Sádlo, J., Bartošová, A., Novák, J., Kovář, P., Jírová, A., Šmilauer, P., Řehounková, K., 2016. Does succession run towards potential natural vegetation? An analysis across seres. J. Veg. Sci. 27, 515–523. 10.1111/jvs.12383

R Core Team, 2010. R: A language and environment for statistical computing.Vienna, Austria: R Foundation for Statistical Computing.Retrieved from http://www.R-project.org/

Roberts, D.W., 2010. labdsv. Ordination and multivariate analysis for ecology.Retrieved from http://cran.r-project.org/web/packages/labdsv/index.html

Ruprecht, E., 2006. Successfully Recovered Grassland: A Promising Example from Romanian Old-Fields. Restor. Ecol. 14, 473–480. 10.1111/j.1526-100X.2006.00155.x

Shannon, C.E., 1948. A mathematical theory of communication. The Bell System Technical Journal 27, 379–423, 623–656.

Silveira, F.A.O., Ordóñez-Parra, C.A., Moura, L.C., Schmidt, I.B., Andersen, A.N., Bond, W., Buisson, E., Durigan, G., Fidelis, A., Oliveira, R.S., Parr, C., Rowland, L., Veldman, J. W., Pennington, R.T., 2022. Biome Awareness Disparity is BAD for tropical ecosystem conservation and restoration. J. Appl. Ecol. 59, 1967–1975. 10.1111/1365-2664.14060

Sonkoly, J., Tóth, E., Balogh, N., Balogh, L., Bartha, D., Bata, K., Bátori, Z., Békefi, N., Botta-Dukát, Z., Bölöni, J., Csecserits, A., Csiky, J., Csontos, P., Dancza, I., Deák, B., Dobolyi, Z.K., E-Vojtkó, A., Gyulai, F., Hábenczyus, A.A., Henn, T., Horváth, F., Höhn, M., Jakab, G., Kelemen, A., Király, G., Kis, S., Kovacsics-Vári, G., Kun, A., Lehoczky, É., Lengyel, A., Lhotsky, B., Löki, V., Lukács, B.A., Matus, G., McIntosh-Buday, A., Mesterházy, A., Miglécz, T. V., A.M., Molnár, Z., Morschhauser, T., Papp, L., Pósa, P., Rédei, T., Schmidt, D., Szmorad, F., Takács, A., Tamás, J., Tiborcz, V., Tölgyesi, C., Tóth, K., Tóthmérész, B., Valkó, O., Virók, V., Wirth, T., Török, P., 2022. PADAPT 1.0 – the Pannonian Database of Plant Traits. bioRxiv, 10.1101/2022.12.05.519136

Sonkoly, J., Valkó, O., Balogh, N., Godó, L., Kelemen, A., Kiss, R., Miglécz, T., Tóth, E., Tóth, K., Tóthmérész, B., Török, P., 2020. Germination response of invasive plants to soil burial depth and litter accumulation is species-specific. J. Veg. Sci. 31, 1079–1087. 10.1111/jvs.12891

Teleki, B., Sonkoly, J., Erdős, L., Tóthmérész, B., Prommer, M., Török, P., 2020. High resistance of plant biodiversity to moderate native woody encroachment in loess steppe grassland fragments. Appl. Veg. Sci. 23, 175–184. 10.1111/avsc.12474

Timár, G., Molnár, G., Székely, B., Biszak, S., Varga, J., Jankó, A., 2006. Digitized maps of the Habsburg Empire – The map sheets of the second military survey and their georeferenced version. Budapest, Hungary: Arcanum.

Tölgyesi, C., Török, P., Kun, R., Csathó, A.I., Bátori, Z., Erdős, L., Vadász, L., 2019. Recovery of species richness lags behind functional recovery in restored grasslands. Land Degr. Develop. 30, 1083–1094. 10.1002/ldr.3295

Tölgyesi, C., Buisson, E., Helm, A., Temperton, V., Török, P., 2022. Urgent need for updating the slogan of global climate actions from “tree planting” to “restore native vegetation”. Restor. Ecol. 30, e13594. 10.1111/rec.13594

Török, P., Vida, E., Deák, B., Lengyel, S., Tóthmérész, B., 2011a. Grassland restoration on former croplands in Europe: an assessment of applicability of techniques and costs. Biodivers. Conserv. 20, 2311–2332. 10.1007/s10531-011-9992-4

Török, P., Kelemen, A., Valkó, O., Deák, B., Lukács, B.A., Tóthmérész, B., 2011b. Lucerne-dominated fields recover native grass diversity without intensive management actions. J. Appl. Ecol. 48, 257–264. 10.1111/j.1365-2664.2010.01903.x

Török, P., Brudvig, L.A., Kollmann, J. N. Price, J., Tóthmérész, B., 2021. The present and future of grassland restoration. Restor. Ecol. 29, e13378. 10.1111/rec.13378

Tóth, E., Deák, B., Valkó, O., Kelemen, A., Miglécz, T., Tóthmérész, B., Török, P., 2018. Livestock type is more crucial than grazing intensity: Traditional cattle and sheep grazing in short-grass steppes. Land Degr. Develop. 29, 231–239.10.1002/ldr.2514

Triantis, K.A., Guilhaumon, F., Whittaker, R.J. (2011): The island species–area relationship: biology and statistics. J. Biogeogr. 39: 215–231. 10.1111/j.1365-2699.2011.02652.x

Turtureanu, P.D., Palpurina, S., Becker, T., Dolnik, C., Ruprecht, E., Sutcliffe, L.M.E., Szabó, A., Dengler, J., 2014. Scale- and taxon-dependent biodiversity patterns of dry grassland vegetation in Transylvania. Agric. Ecosys. Environ. 182, 15–24. 10.1016/j.agee.2013.10.028

United Nations, 2019. UN Decade of Ecosystem Restoration 2021-2030, resolution A/RES/73/284on 1stMarch 2019.

Veldman, J.W., Buisson, E., Durigan, G., Fernandes, G.W., Le Stradic, S., Mahy, G., Negreiros, D., Overbeck, G.E., Veldman, R.G., Zaloumis, N.P., Putz, F.E., Bond, W.J., 2015. Toward an old-growth concept for grasslands, savannas, and woodlands. Frontiers Ecol. Environ. 13, 154–162. 10.1890/140270

Villéger, S., Mason, NW., Mouillot, D., 2008. New multidimensional functional diversity indices for a multifaceted framework in functional ecology. Ecology 89, 2290–2301. 10.1890/07-1206.1

Virágh, K., Bartha, S., 2003. Species turnover as a function of vegetation pattern. Tiscia. 34, 47–56.

Wesche, K., Ambarli, D., Török, P., Kamp, J., Treiber, J., Dengler, J., 2016. Thepalaearctic steppe biome: a new synthesis. Biodivers. Conserv. 25, 2197–2231. 10.1007/s10531-016-1214-7

Wilson, J.B., Peet, R.K., Dengler, J., Pärtel, M., 2012. Plant species richness: the world records. J. Veg. Sci. 23, 796–802. 10.1111/j.1654-1103.2012.01400.x

Zhang, J., Gillet, F., Bartha, S., Alatalo, J.M., Biurrun, I., Dembicz, I., Grytnes, J.-A., Jaunatre, R., Pielech, R., Van Meerbeek, K., Vynokurov, D., Widmer, S., Aleksanyan, A., Bhatta, K.P., Campos, J.A., Czortek, P., Dolezal, J., Essl, F., García-Mijangos, I., Guarino, R., Güler, B., Hájek, M., Kuzemko, A., Li, F.Y., Löbel, S., Moradi, H., Naqinezhad, A., Silva, V., Šmerdová, E., Sonkoly, J., Stifter, S., Talebi, A., Török, P., White, H., Wu, J., Dengler, J., 2021. Scale dependence of species–area relationships is widespread but generally weak in Palaearctic grasslands. J. Veg. Sci. 32, e13044. 10.1111/jvs.13044

